# Robust CNV detection using single-cell ATAC-seq

**DOI:** 10.1101/2023.10.04.560975

**Authors:** Travis W. Moore, Galip Gürkan Yardımcı

## Abstract

Copy number variation (CNV) is a widely studied type of structural variation seen in the genomes of cancerous and other dysfunctional cells. CNVs can have direct and indirect effects on gene dosage, and are thought to drive cancer progression and other disorders. Advancements in single-cell assays such as sc-ATAC-seq and sc-RNA-seq, along with their ubiquitous use, allows for the identification of CNVs at single cell resolution. While there are a variety of available tools for CNV detection in sc-RNA-seq, development of sc-ATAC-seq based accurate and reliable CNV callers is in the early stages, with only two available algorithms so far. We present RIDDLER, a single-cell ATAC-seq CNV detection algorithm based on outlier aware generalized linear modeling. By utilizing tools from robust statistics, we developed an extensible model that is able to identify single-cell CNVs from sc-ATAC-seq data in an unsupervised fashion, while providing probabilistic justification for results. Our statistical approach also allows us to estimate when loss of signal is likely caused by drop-out or a true genome deletion event, as well as predict reliable CNVs without the need for normative reference cells. We demonstrate the effectiveness of our algorithm on cancer cell line models where it achieves better agreement with bulk WGS derived CNVs than competing methods. We also compare our approach on 10x multimone data, where it shows better agreement and integration with RNA derived CNV estimates.

## 1 Introduction

Copy number variations (CNV) are one of most well studied types of genetic structural variation. Broadly speaking, CNVs correspond to the phenomenon where a particular region of a genome has fewer or more than two copies, in the case of a diploid genome. CNVs can dramatically vary in size, ranging from entire chromosomes to hundreds of kilobases. Naturally, CNVs can change gene dosage due to gain or loss of copies[1], or have direct or indirect effects on gene regulation since non-coding regulatory regions may be lost or misplaced due to their effects. There are a variety of development disorders and diseases that have been linked to CNVs[2, 3, 4], and due to their potential for impacting gene regulation, they may impact the state of healthy cell populations in a subtle and heterogeneous manner. The prevalence of CNVs in cancerous cells have been deeply documented and is thought to play a critical role in cancer development and progression in many cancer types. Thus, fast and accurate detection of CNVs is valuable for researchers studying cancer, development disorders, and links between gene regulation and genomic structural variations.

Contemporary approaches for CNV detection heavily utilize next-generation sequencing data-sets due to the prevalence and standardization of these assays. Whole genome sequencing (WGS) and exome sequencing naturally lend themselves to detection of CNVs; these assays uniformly cover the whole genome or the exome, after accounting for assay specific biases. Thus, a loss or a gain of a genomic region is clearly reflected as a decrease or increase of coverage compared to the baseline coverage of the genome. Additionally, recently research has reported successfully utilizing genomics assays that measure transcriptome and three dimensional genome folding -bulk tissue RNA-seq[5, 6, 7] and Hi-C[8, 9, 10] - for CNV detection. With the proliferation of data-sets resulting from such assays and their broad availability, CNV detection algorithms that can be applied to a variety of genomics assays are valuable for both better utilization of these assays for CNV detection beyond their original purpose, and linking CNVs to transcriptomic and epigenetic states. Unfortunately, the bulk tissue nature of these assays result in averaging of CNV profiles across many individual cells, making it challenging to study CNVs that might be manifest in a subset of cells within the bulk cell population. An obvious example is the heterogeneous nature of cancerous cell populations, consisting of multiple clones.

The recent explosion of single-cell genomics assays raises the possibility of utilizing these data-sets to detect CNVs at single-cell resolution, counteracting the weaknesses of population averaging effects for bulk genomics assays. Indeed, there are multiple algorithms to detect CNVs at single-cell resolution from sc-RNA-seq assay, such as inferCNV[11], HoneyBadger[12], CopyKAT[13], and SCEVAN[14]. sc-RNA-seq data-sets have been utilized by many research groups and consortia to profile the transcriptome of many different heterogeneous cell populations, tissues, and even whole biological systems, making them a great resource for CNV detection. However, sc-RNA-seq based CNV callers are relatively challenged by the uneven coverage of coding sequences along the genome, and may require reference cell populations to detect CNVs.

In this study, we introduce Robust Inference of Duplications and Deletions using LinEar Regression (RIDDLER): a novel CNV calling algorithm that utilizes arguably the second most prevalent single-cell genomics assay, sc-ATAC-seq. sc-ATAC-seq measures the accessibility of chromatin in genome wide fashion at single-cell resolution, detecting both coding and non-coding regions of the genome. Our algorithm utilizes robust statistics and a generalized linear model to detect CNV gains and losses in an unsupervised fashion without the need for additional reference cells. We compared our approach against recently published methods Copy-scAT[15] and EpiAneufinder[16] on public sc-ATAC-seq data from cancer cell lines. In these experiments our approach consistently achieved a better balance between comprehensive detection of true CNV events and avoidance of false positives, as measured by sensitivity and specificity. To show the usefulness and limitations of sc-ATAC-seq based CNV detection, we thoroughly characterize the accuracy of RIDDLER under varying levels of sparsity and describe the necessary levels of ATAC-seq reads per cells to detect the majority of CNV gain and loss events. We show that RIDDLER can operate at different resolutions ranging from 1MB to 100kb, and can detect CNVs at much higher resolution compared to Copy-scAT. Lastly, we demonstrate a powerful application of RIDDLER for CNV based integration of single-cell populations profiled by sc-ATAC-seq and sc-RNA-seq using sc-multiomics assays applied to two different breast cancer cell lines. These results demonstrate the superior performance and versatility of our generalizable CNV detection algorithm for robust detection of CNVs at single-cell resolution.

## 2 Results

### 2.1 Outlier-aware CNV detector for scATAC-seq using robust statistics

Our CNV detection algorithm is built on the idea of computing the expected signal across the genome via a predictive model, and then finding significant deviations from that expected signal within the observed data. Our predictive model utilizes a robust generalized linear model (GLM)[17] with a Poisson distribution. Using a robust GLM, optimized via Huber loss, is critical to our approach as it allows us to effectively ignore the CNV driven signal. Specifically, Huber loss down-weights points with high absolute residuals, reducing the influence of outliers when fitting the model. We expect the majority of genome regions across cells in a sample to be diploid, with fewer non-diploid CNV regions mixed in. Thus a robust model should reduce the influence of CNVs and provide a better estimate of diploid behavior; indeed Supplemental Figure 1 shows that a non-robust version of our GLM has dramatically reduced performance. Finding deviations from an expected distribution is a common approach used in many CNV callers[18, 13, 11, 15], often with a reference dataset or additional inputs used to define the distribution corresponding to diploid cells. By utilizing robust GLMs, we are able to model the diploid distribution within the target sample, eliminating the need for a separate reference population.

Identifying CNVs from single cell assays is difficult due to well known challenges, such as data sparsity and dropout. Along with our robust modeling framework, we also implement a processing pipeline to better account for these challenges. Our pipeline begins by binning scATAC-seq reads into regularly spaced windows across the genome, creating a window by cell matrix of fragment counts which is then normalized by read depth. The sizes of these windows can be set based on the average read depth of the data, with deeper sequenced data able to accommodate finer resolution windows. The genome covariates of the model (GC content, mappability, peak coverage) are also averaged over the same sized windows. We run this data matrix through a drop-out detector, which uses a simple sampling distribution model to identify zero-read matrix values that are likely due to signal drop out. These drop-out windows are removed from downstream analysis to prevent them being called as CNV losses. Once our robust GLM model is fit and the residuals of each window in each cell calculated, we use a sliding window p-value calculation to identify outliers that are statistically significant as CNVs. The sliding window approach is used to smooth out variations within individual cells, making CNVs more likely to be called if they are adjacent to other regions of similar deviation from the expected signal. CNVs that are deemed statistically significant are then assigned a copy number based on the magnitude of their difference from the expected distribution.

We use measurements of GC content, mappability, and peak coverage across the genome as the covariate inputs to the GLM for scATAC-seq. GC content and mappability are are commonly used covariates when accounting for coverage biases that should be universal across the assay. Peak coverage is the only ATAC-seq specific covariate, which aims to capture the fluctuating distribution of ATAC-seq peaks along the genome. A window that is covered by more peaks will emit more ATAC-seq signal and vice versa. We use peaks from a multi-tissue reference set[19] for all experiments, which allows a universal option for this input. In theory, RIDDLER can be extended to other modalities by using alternate covariates that account for biases in those assays, as further elaborated in the Discussion section.

### 2.2 RIDDLER accurately detects single-cell CNVs

To evaluate the ability of RIDDLER to detect CNVs at single cell resolution, we tested the algorithm on single-cell ATAC-seq data generated from the gastric cancer cell line SNU601[18]. This cell line has previously been shown to contain both large scale gains and losses spanning several megabases in size. To act as ground truth for comparison, we used scWGS of the same cell line[20]. The WGS reads were bulked and passed to HMMcopy[21], run with a 6 state model. We bulk the signal to ensure this ground truth has the highest possible data quality, and with the assumption that a cultured cancer cell line should show minimal heterogeneity.

We compared the performance of RIDDLER to other scATAC-seq CNV callers Copy-scAT[15] and epiAneufinder[16].Both competitor algorithms were run with the default suggested parameters. RIDDLER, epiAneufinder, and HMM-copy all used the same 1MB window size for data, while Copy-scAT can only report calls at chromosome arm resolution. We split the CNV calls of Copy-scAT to 1MB windows so that it can be compared to HMMcopy at the same resolution as all other algorithms.

For each scATAC-seq algorithm, we compare the CNV calls at each window of each cell to the same window of the WGS calls. All CNVs were converted to the three state output of loss, normal, gain, to simplify this comparison. We report the concordance of scATAC calls with WGS calls using measures of F1-score, sensitivity, and specificity. These are computed for the target classes of loss and gain separately (panels E and F of Figure 2), as well as both combined (panel D). The distributions of these scores across all cells are shown as violin plots. We also add a single point to these plots, representing the score of the consensus CNV calls for each algorithm across all cells. This consensus call is calculated across each window as all CNVs that appear in at least 25 percent of cells, or the majority call if both gains and losses are called in the same window. We expect cell lines to have minimal heterogeneity between cells, so these consensus calls represent the strongest and most consistent signal from each caller. These consensus scores are shown as black dots on the violin plots.

**Figure 1:**
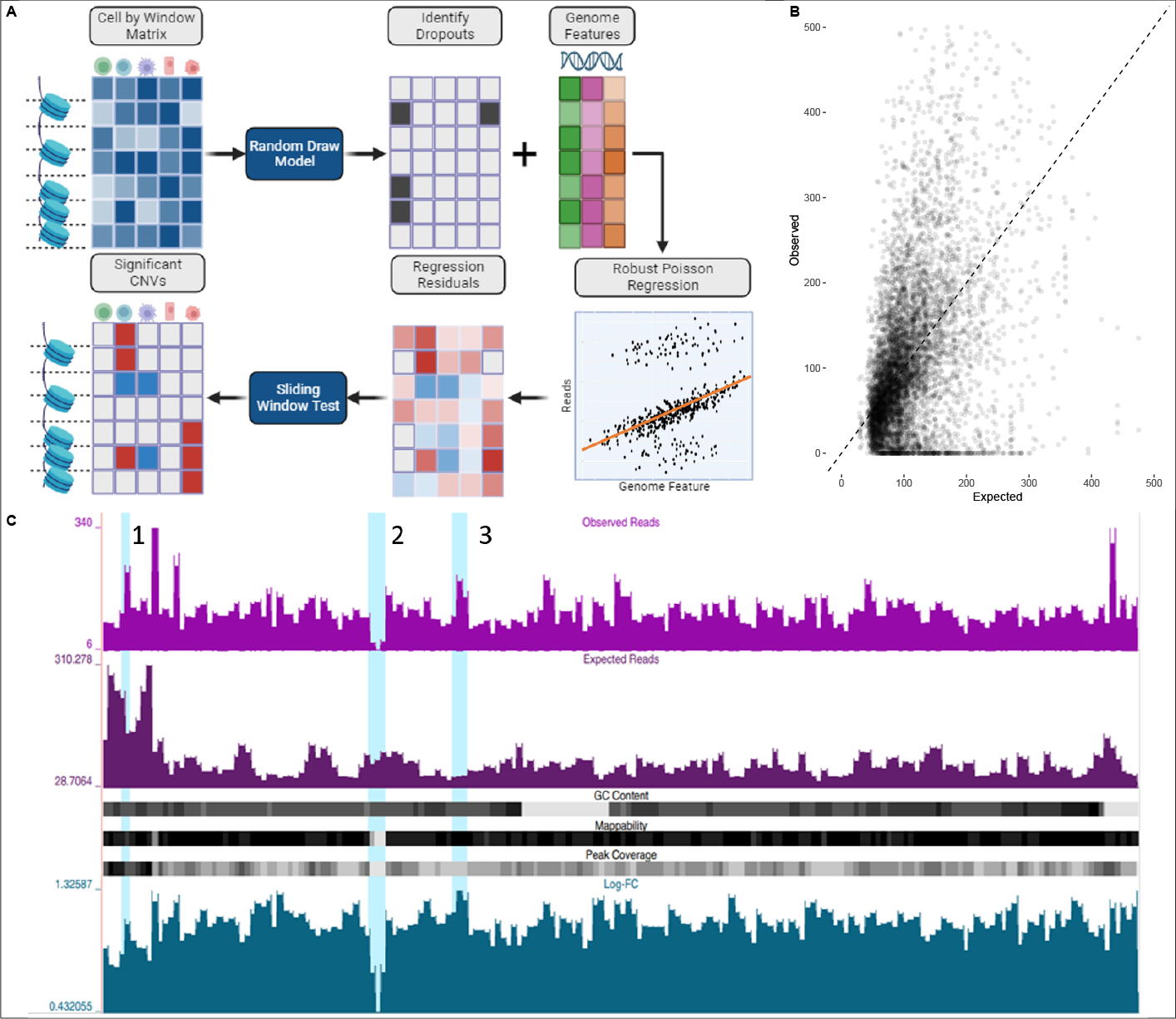
A) RIDDLER pipeline. Reads are binned per cell into fixed length windows to create the input matrix, which is normalized and filtered for drop-outs. A robust Poisson regression model is fit across all cells along with genomic features for each window. A sliding window test is then applied across the model residuals to identify significant CNVs and assign copy numbers. B) Example distribution of observed reads from normalized data and expected reads from model fitting, from SNU601 cell. C) Genome browser view of observed reads, expected reads, and genome features for chromosome 4 of SNU601 cell. Region 1 highlights where an increase in observed signal is explained by increases in the features, while 2 and 3 highlight potential loss and gain areas, respectively.

**Figure 2:**
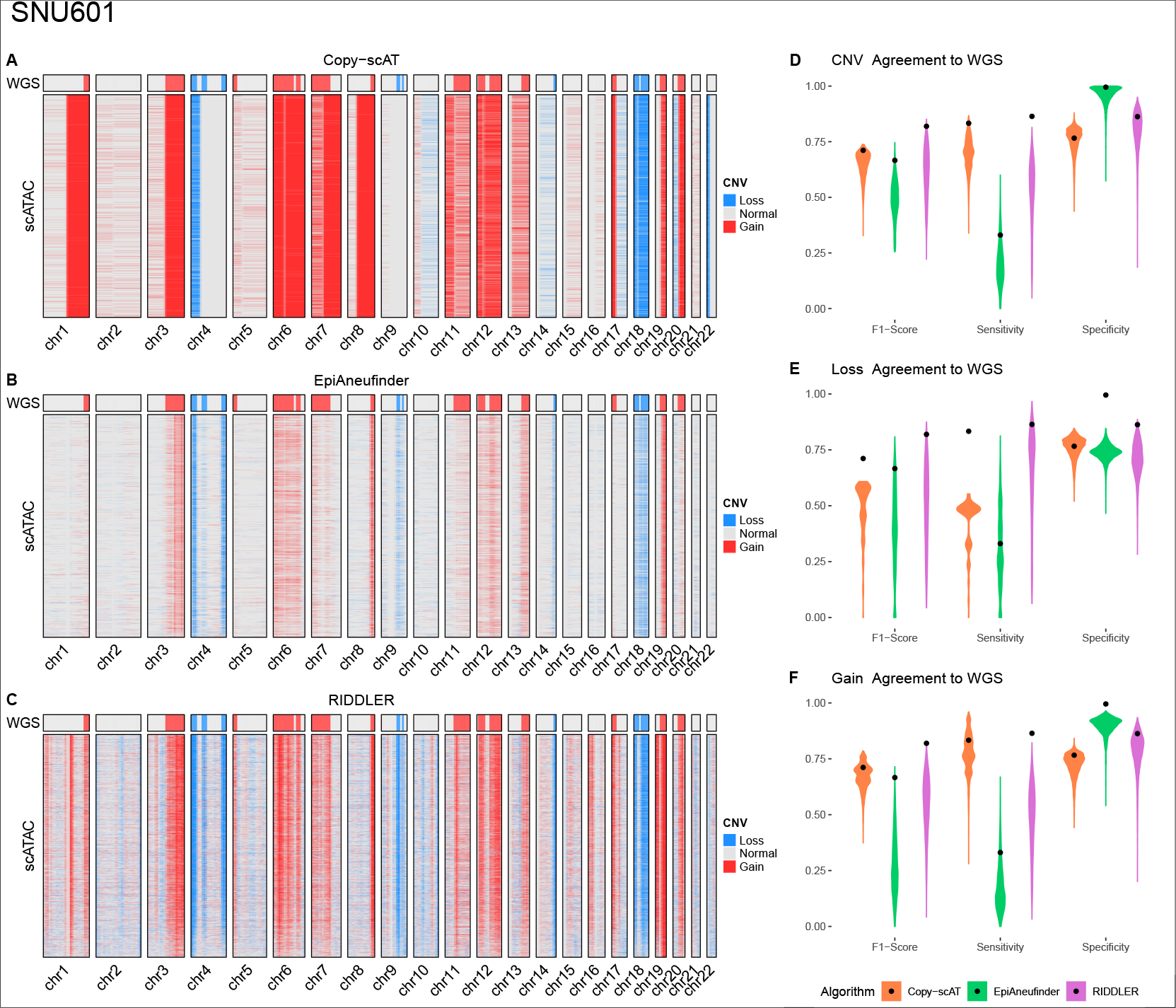
A-C) Heatmaps of CNV calls for each algorithm on SNU601 cell line, with WGS CNVs as reference. D-F) Metrics comparing predicted CNVs to WGS calls. Metrics are shown as a distribution across each cell, with a black dot indicating the value for the consensus calls across all cells.

Figure 2 shows that all three scATAC callers identify CNVs that are validated by WGS. Copy-scAT has very consistent calls for most CNVs, but lacks the resolution to call gains or losses at sub-arm levels. EpiAneufinder makes sparser, more conservative calls, resulting in very high specificity and low sensitivity. RIDDLER strikes a balance between these two methods, achieving good sensitivity and specificity, resulting in an improved F1-score. RIDDLER also does consistently well in identifying both gains and losses, compared to Copy-scAT which misses the smaller resolution losses, and epiAneufinder which misses the more subtle gain events.

### 2.3 RIDDLER maintains high specificity with reduced signal and increasing resolution

A natural extension to RIDDLER is to identify CNVs at finer resolutions, achieved by reducing the bin size for windows. We investigated the effects of running RIDDLER on the SNU601 cell line data with window sizes of 1MB, 500KB, 250KB, and 100KB. Figure 3 shows that the F1-scores for the algorithm remain relatively consistent up to 250KB, with a more noticeable drop-off at 100KB resolution. This reduction in performance is due entirely to a decline in the sensitivity of the algorithm, as the same CNVs are called less consistently with smaller window sizes. This is expected as only cells with the highest coverage will be able to maintain predictive performance with smaller windows. Notably the specificity of the algorithm remains consistent during these experiments.

**Figure 3:**
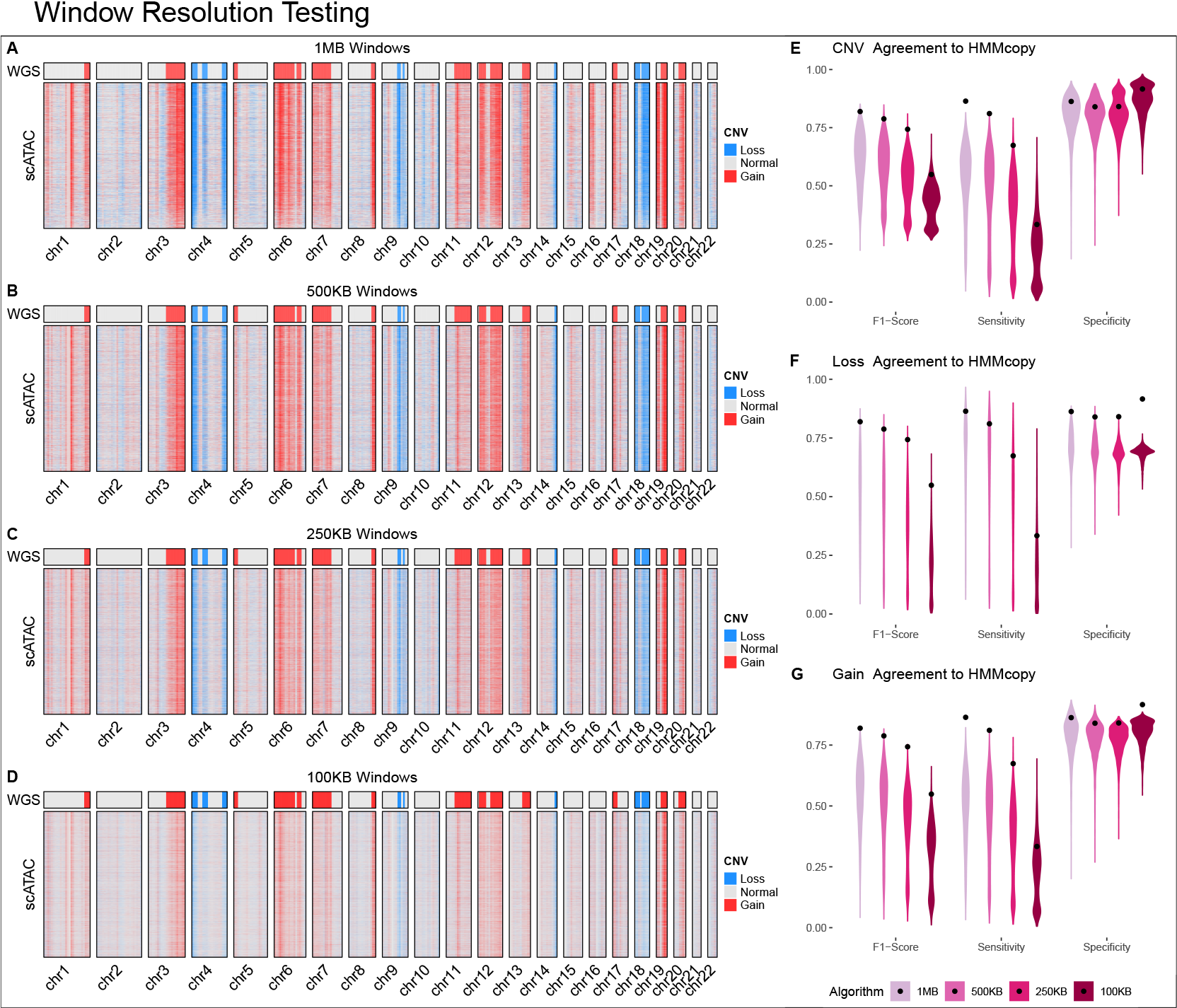
A-D) Heatmaps of CNVs reported by RIDDLER on SNU601 data with window size set to 1MB, 500KB, 250KB, and 100KB. E-G) Metrics comparing predicted CNVs to WGS calls. Metrics are shown as a distribution across each cell, with a black dot indicating the value for the consensus calls across all cells.

Reducing the window size reduces the average number of reads per window, diluting the overall signal of the data. We further examined the performance of RIDDLER with reduced signal by holding the window size fixed at 1MB and down-sampling the total reads in each cell. We did this for read depths of 20,000, 15,000, 10,000, and 5,000, as shown in Figure 4. The original scATAC-seq data had very high coverage, cited as a mean of 73,845 fragments per cell, so there is an expected drop off in performance with sparser cells. Just like with the window reduction experiments, the specificity of the algorithm remains consistent while the sensitivity slowly declines, with a sharper drop off at a read depth of 10,000. This indicates that the algorithm maintains good performance up to an average of approximately 5 reads per window.

**Figure 4:**
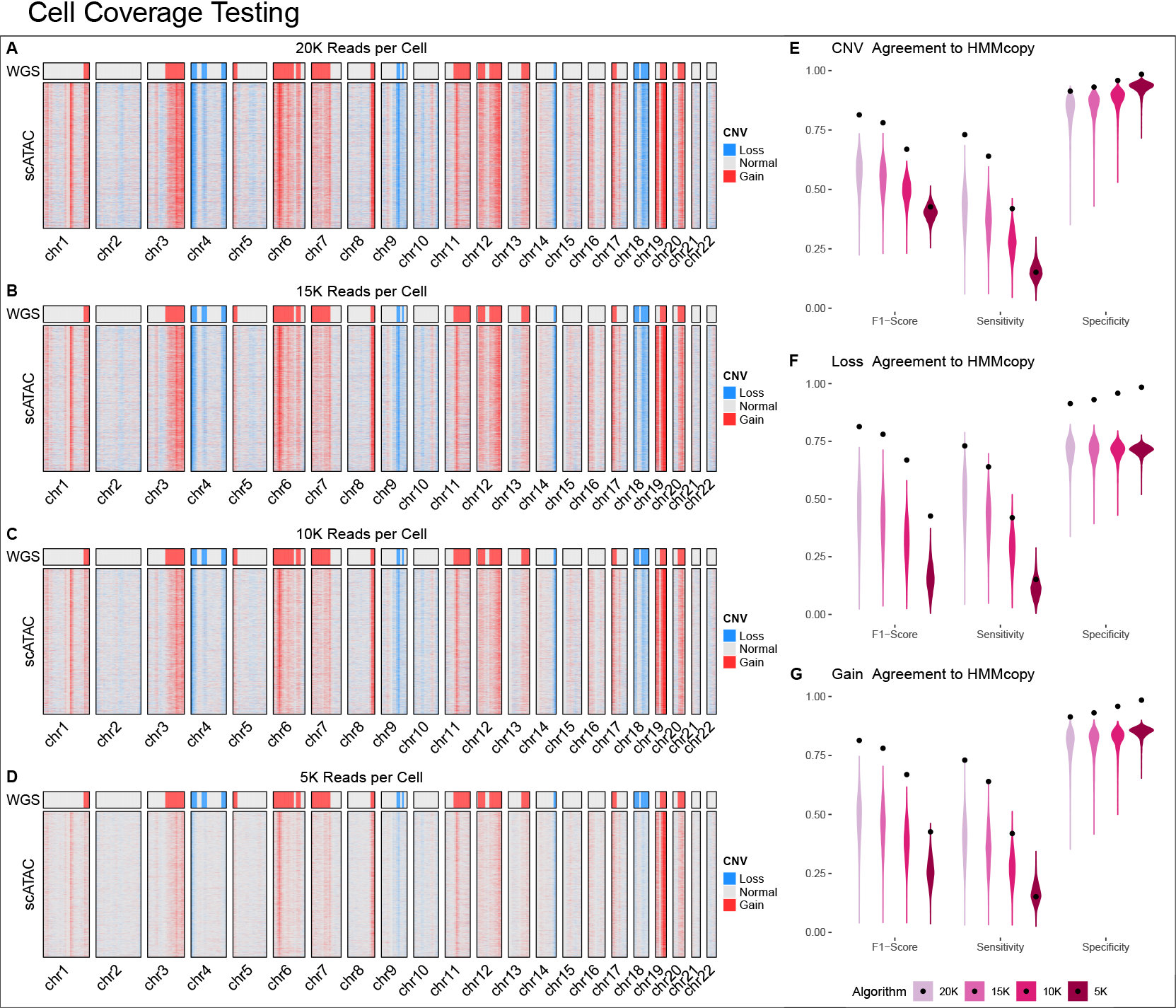
A-D) Heatmaps of CNVs reported by RIDDLER on SNU601 data with each cell down-sampled to 20, 15, 10, and 5 thousand reads. E-G) Metrics comparing predicted CNVs to WGS calls. Metrics are shown as a distribution across each cell, with a black dot indicating the value for the consensus calls across all cells.

While a decline in performance is unavoidable as the signal per window becomes more diluted, we emphasize that the calls RIDDLER does make remain trustworthy. Our robust regression approach reliably models normative signal, ensuring that only regions that are statistically distinct from the expected signal are called as CNVs. While a reduction in signal makes these deviations more difficult to find consistently, it does not result in the hallucination of new CNVs. In particular, our dropout filter helps to ensure that spurious losses are not called when signal poor regions become more frequent.

### 2.4 RIDDLER reliably identifies CNVs in single-cell multiomics assays

We evaluated RIDDLER and other scATAC callers on 10x Multiome data generated from breast cancer cell lines T47D and MCF7[22]. The 10x Multiome assay is a single-cell assay which yields both sc-RNA-seq and sc-ATAC-seq readout from the same cell. Thus we have both sc-ATAC-seq and sc-RNA-seq readout of the same MCF7 and T47D cells. For both of these datasets, we have lower ATAC read counts per cell than the SNU601 data, with 16,000 and 11,000 median reads for T47D and MCF7 respectively, making them a more difficult dataset to work with. In the absence of matching WGS data from these cell lines, we instead used inferCNV[11] to compute CNVs from the paired scRNA for use as an imperfect ground truth evaluation. As shown in Supplemental Figure 3, inferCNV did not show any signs of subclonal populations within either cell line. Given this, and the expectation of minimal heterogeneity within cell lines, we computed consensus CNVs for the RNA modality to use as a single ground truth evaluation for each ATAC cell CNV.

Figure 5 shows the results of each scATAC caller compared to consensus RNA calls. Again we see a similar trend where copy-scAT has high sensitivity but low specificity, epiAneufinder has low sensitivity with high specificity, and RIDDLER strikes a balance between the two with slightly higher F1-score. In general both of these cell lines contain a large amount of relatively small gain and loss events, as reported by inferCNV, making them particularly challenging for copy-scAT with its limited resolution. Copy-scAT also reports distinct heterogeneity in the cell lines, switching between gain and loss in chromosome 18 of T47D and chromosome 6 of MCF7 (Supplemental Figure 2), which does not match the homogeneous RNA calls. EpiAneufinder and RIDDLER make many of the same calls, but RIDDLER is more consistent with its calls across all cells. Overall, we see less concordance with the RNA CNVs than with the WGS CNVs of SNU601, though as noted RNA is an imperfect ground truth.

**Figure 5:**
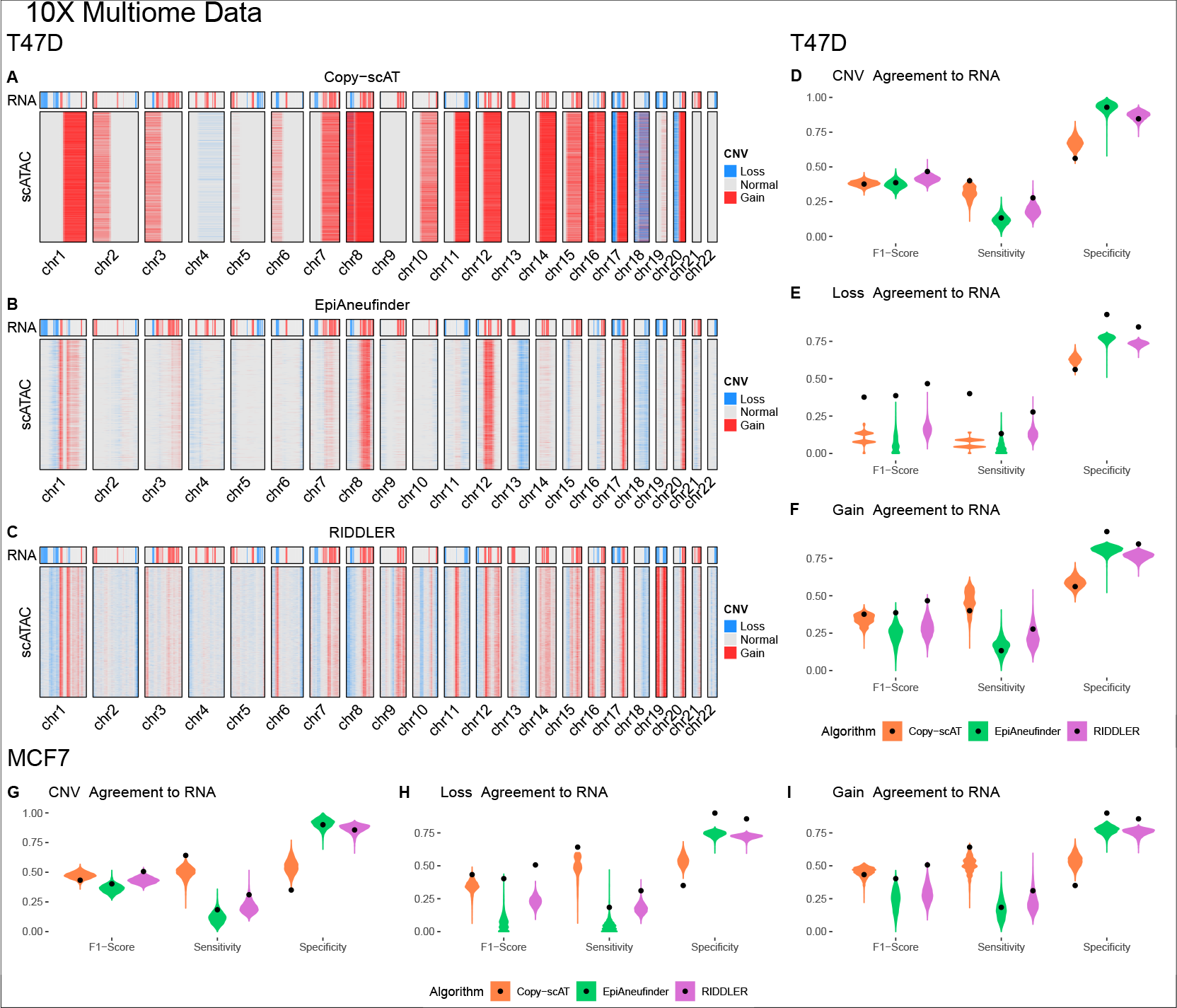
A-C) Heatmaps of CNV calls for each algorithm on T47D cell line, with RNA CNVs as reference. D-F) Metrics comparing predicted CNVs to RNA calls for T47D cell line. G-I) Metrics comparing predicted CNVs to RNA calls for MCF7 cell line.

### 2.5 RIDDLER accurately integrates single-cells based on RNA and ATAC derived CNV profiles

As an additional evaluation of scATAC CNV calls for the 10x data, we performed an integration experiment based on CNV profile similarity. We mixed the T47D and MCF7 datasets in-silico, creating a two cell line dataset with both ATAC and RNA readouts for each cell. For each cell in the combined dataset we used the ATAC based CNV profile to find the cell with the best matching RNA based CNV profile from inferCNV. We computed the best match between RNA based and ATAC based CNV profiles using euclidean distance, with loss represented as -1, normal as 0, and gain as 1. After the best match was computed for each cell, we tracked how often cells from the same cell line were matched together when using the predictions of each ATAC caller. Multiple studies have performed integration of sc-RNA-seq and sc-ATAC-seq datasets based on a transcriptomic axis[23, 24, 25]; our goal in this experiment is to demonstrate the potential of a CNV based integration between the two modalities.

Ideally, we expect the majority of cells in the combined dataset to match to cells of the same type after integration. Figure 6 shows the results for each ATAC caller, with CNV profiles identified by RIDDLER having a significantly higher rate of correctly matching RNA CNVs from the same cell types when compared to CNV profiles identified by copy-scAT and epiAneufinder. The CNV UMAPs also show that both Copy-scAT and epiAneufinder have distinct subgroups within each cell line, while inferCNV and RIDDLER have a single cluster for each cell line. This indicates that the between cell CNV variability for these two methods was closer to white noise than consistent structural differences, which would be expected with lab cultured cell lines.

**Figure 6:**
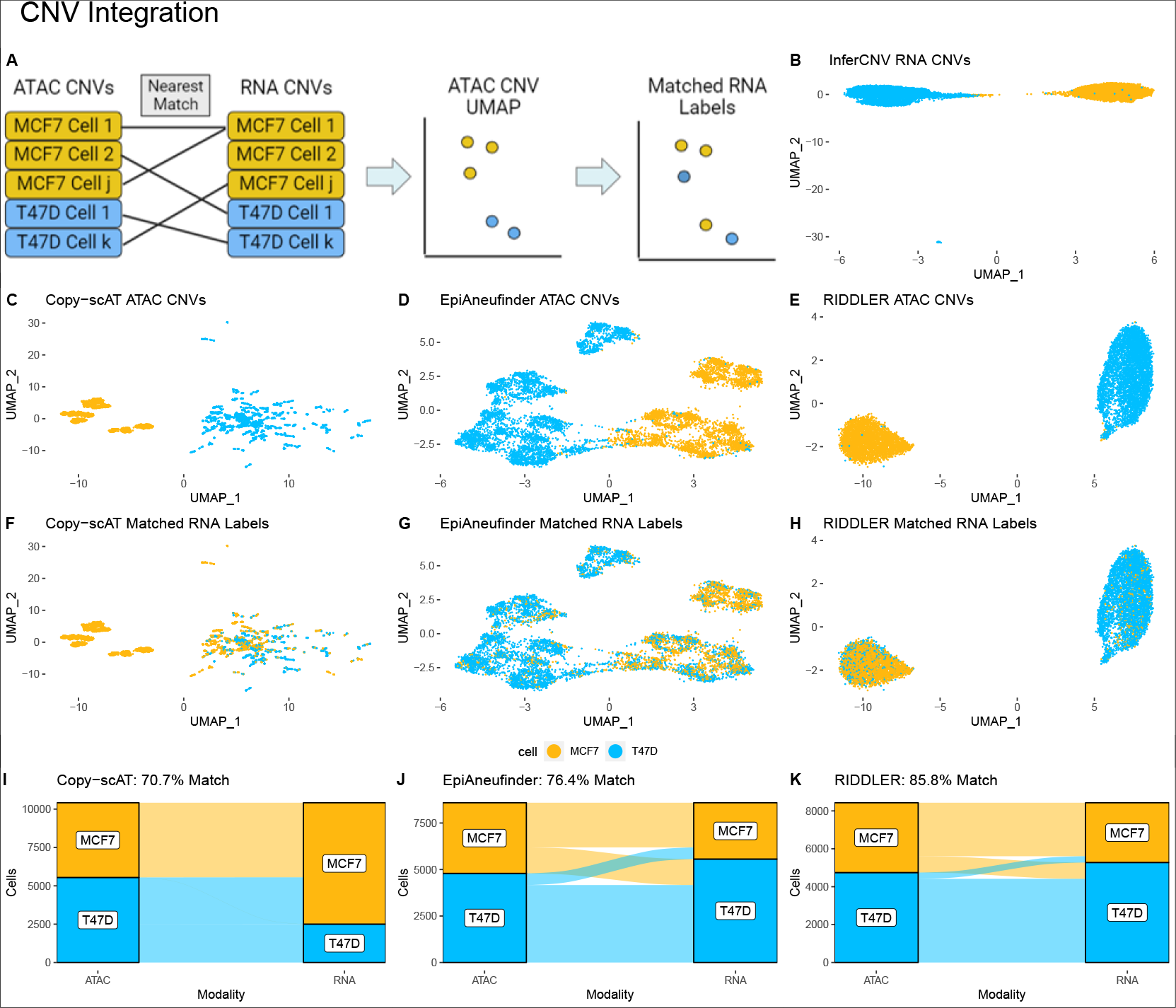
A) Schematic of integration procedure. B-E) UMAPs of CNV calls for each algorithm. F-H) UMAPs of ATAC calls, relabeled with matched RNA cell types. I-K) Alluvial plots of ATAC/RNA matching for each ATAC caller.

## 3 Discussion

We demonstrated that RIDDLER can accurately detect CNV events within a single cell using sc-ATAC-seq data in an unsupervised fashion, while maintaining most of its ability to call CNVs at higher resolutions and increasing levels of data sparsity. We thoroughly benchmarked RIDDLER and two competing methods’ ability to identify CNVs at single-cell resolution, using sc-ATAC-seq and 10x Multiome datasets from different cancer cell lines. To validate our CNV calls, we used two strategies: (1) using CNVs identified from matching bulk WGS data, as this is the most widely used approach in the field, (2) using CNVs identified from sc-RNA-seq from the same cells when using 10x Multiome assays. Our results clearly show RIDDLER outperforms competing methods as measured by the F-score, which is the harmonic mean of precision and recall. Additionally, we found that epiAneufinder consistently has high specificity but low sensitivity, and copy-scAT shows the opposite trend. Copy-scAT has the additional limitation of only calling CNVs at arm resolution, which is likely the cause of its high specificity at the cost of low sensitivity. Taken together, RIDDLER strikes a better balance between identifying most of the gold standard CNVs while minimizing false discoveries. Furthermore, consensus of single-cell CNV calls identified by RIDDLER from cancer cell lines exhibit the highest concordance to bulk WGS calls, showing that RIDDLER consistently identifies the same gain and loss events across multiple single-cells, as expected when applied to cells from relatively homogeneous cancer cell lines.

A common challenge in the development and application of computational methods that utilize single-cell assays is data sparsity, especially for approaches that require characterization and identification of fine-scale genomic events. While recent advancements have resulted in significant increments to the number of fragments per cell in a sc-ATAC-seq experiment, numbers still vary significantly across different datasets. Our in-depth analysis of RIDDLER’s performance when applied to sparser and sparser cells shows that RIDDLER does not identify false positives in sparser cells, though increasing sparsity does lead to an expected loss in sensitivity. We attribute some of this robustness to our drop-out filter, which statistically identifies zero-read windows that are likely due to lack of signal. This filter prevents spurious losses from being called as the number of fragments is reduced. It is encouraging to see that performance of RIDDLER doesn’t dramatically suffer until signal is reduced to five thousand fragments per cell, while it maintains most of the CNV calls at ten thousands fragments.

Calling CNVs at single-cell resolution holds great promise in studying heterogeneous cell populations with different underlying CNVs profiles; an obvious application is identification of different clones in a tumour or a cancerous cell population. Unfortunately, the data-sets in this study are from cancer cell lines without a presumed clonal heterogeneity. However, the experiments we ran when comparing and mixing MCF7 and T47D 10x Multiome data revealed very promising results. We showed that RIDDLER identifies different CNV profiles across the two cell lines, thus showing that it can distinguish between cell lines with potentially different CNV profiles at single-cell resolution. Moreover, RIDDLER and inferCNV both report homogeneous CNV profiles within MCF7 and T47D single cell populations according to the UMAP embeddings of both cell lines in Figure 6, as one would expect from cancer cell lines. UMAP embeddings of these cells based on epiAnuefinder and CopyscAT derived CNV profiles seem to identify multiple sub populations of cells with different CNV profiles. Strikingly, matching of T47D and MCF7 cells based on their RNA based CNV profiles from inferCNV and ATAC based CNV profiles from RIDDLER and other competitors revealed that RIDDLER achieves the most accurate matching of cell lines. These results, taken together, show that RIDDLER holds great promise in studying heterogeneous cancerous and other cell populations with different CNV profiles, and integration of same cell populations studied by different single-cell assays on a CNV based axis. Indeed, even within the single-ATAC-seq modality, one can identify CNVs accurately using RIDDLER and correlate CNV profiles to changes in chromatin accessible regulatory landscape and transcription factor binding events via motif enrichment and footprinting.

RIDDLER holds great promise for extension to other single-cell omics assays. Our simple yet intuitive formulation of the linear model - where observed signal in a given genomic window is fit to assay-appropriate covariates, together with the Huber loss function that down weights the CNV driven signal changes in an unsupervised fashion - makes it straightforward to extend our algorithm to other assays. Indeed, we recently adjusted RIDDLER for application to sc-WGS-seq data with a relatively simple change of appropriate covariates; this application was also used to identify CNV rich and CNV depleted cell populations. These results are set to appear in another study that is in press. We anticipate successful adaptation of RIDDLER to other single-cell assay modalities and potentially to sc-multiomics assays, such as the 10x Multiome. With such advancements, we can utilize our algorithm for both high confidence CNVs validated by different modalities, and also aspire to identify multiscale and high resolution CNVs via utilization of data rich multimodal sc-multiomics datasets. Indeed, future applications of RIDDLER to such high confidence and high resolution CNV calls from sc-multiomics data holds great potential in linking CNV driven phenotype changes at single-cell resolution, getting as close as possible to causal linking of CNVs to cell state.

## 4 Methods

### 4.1 Data Availability

The scATAC-seq and scWGS datasets for the SNU601 cell lines were obtained from the Sequence Read Archive (SRA), accession numbers PRJNA674903 and PRJNA498809, respectively.

10x Multiome data of paired scATAC-seq and scRNA-seq for T47D and MCF7 cell lines were obtained from the Gene Expression Omnibus (GEO) database with accession number GSE154873.

PBMC 10x scATAC-seq data was downloaded from the 10x genomics website https://support.10xgenomics.com/single-cell-multiome-atac-gex/datasets. The dataset title was ‘PBMC from a Healthy Donor - Granulocytes Removed Through Cell Sorting (10k)’. This PBMC dataset was used as the reference cell set when running inferCNV on both the MCF7 and T47D cell lines.

### 4.2 Codebase

Code for running RIDDLER can found on Github, https://github.com/yardimcilab/RIDDLER.

### 4.3 Robust-CNV Pipeline

#### 4.3.1 Normalization

To account for differences in read depth, each cell in a sample is normalized by its trimmed mean. Using the binned number of reads for all windows of a cell, we trim the top and bottom 15th percentile of values to remove outliers, and compute the average of the remaining windows. All windows in the cell are divided by this trimmed mean value, and then multiplied by 100 and rounded to the nearest whole number. We perform this operation on each cell individually.

#### 4.3.2 Drop-Out Filtering

A common issue in single cell analysis is dealing with data sparsity. Put another way, it is often difficult to determine if a cell region with zero reads is due to technical errors, or a true biological loss of signal (such as a chromosome deletion). We use a probabilistic estimation to identify which zero values in the data are likely due to biological events, and which are likely due to sparsity.

Intuitively, when zero values occur in high probability windows for cells with many reads, we have a higher confidence that they are due to true biological events. Conversely, when they occur at low probability sites in cells with few total reads, they are more likely due to sparsity. In our approach, we estimate the average probability of a read occurring in each window, and then calculate the likelihood of no reads falling in a given window for a cell with *n* total reads.

Using the entire input matrix *M* of reads per cell (*i*) per window (*j*), we compute the probability of a read randomly being drawn for each window.

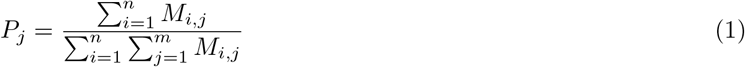

We make the simplifying assumption that these probabilities are the same for each cell. Next, we compute the likelihood of a window having zero reads, assuming reads are distributed randomly according to the probability of each window.

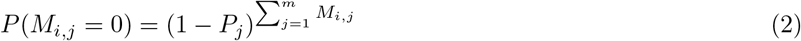

These probabilities give us the likelihood that a zero value would occur by random chance, given the average frequency of the region and the coverage of the cell. We use an FDR correction of these probabilities (*α* = 0.01) over the entire matrix to select a cutoff probability for zero values. Any zero values with probabilities below this threshold are treated as true zeros, while those above the probability are removed from analysis.

#### 4.3.3 Robust GLM Training and Prediction

We use the robustbase package in R to fit a robust Poisson GLM, which implements the method described in Cantoni and Ronchetti[17]. The response values for the GLM are a vectorized version of the normalized cell by window matrix values, denoted as *Y*, with those tagged as drop-outs removed. For each input value *Y*_*i*_, we include the covariates of the corresponding window, denoted as *X*_*i*_, creating a feature matrix *X*. Rows of *X* only depend on window location, with every cell having the same feature values for the same windows. The GLM is fit on the formula *Y∼X*, with an intercept term included.

Once the model has been trained, we compute expected values by predicting the response for each window, given that window’s covariates. Since the covariates only vary by window locations, each cell will have the same vector of expected values across windows.

#### 4.3.4 Genomic features for ATAC-seq

GC content was computed using the bedtools nuc command for the given window resolution.

Mappability scores for a window were computed using GEM[26]. The resulting bigwig file was averaged over bin windows using the Encode software function bigWigAverageOverBed (https://www.encodeproject.org/software/bigwigaverageoverbed/).

Peak coverage per window was computed with respect to a multi-tissue peak reference set of DNase I hyper-sensitive sites[19]. We used the full set of locations for hg38, downloaded from https://zenodo.org/record/3838751 as an all purpose peak set for each experiment. Peak coverage was then computed as the total length of peaks overlapping each window divided by the window size.

In general the peak set used for RIDDLER can be generated from the ATAC data used as input to the model. However, if there is a lack of diversity in cell types within the data, as there was with the different cell lines we tested in this paper, then the peak set will be too predictive of signal and make CNVs more difficult to identify. An ideal use-case would be to use a peak set derived from diploid cells of the same type as those being queried, though as we’ve shown the multi-tissue set is sufficient to capture common accessibility trends.

### 4.3.5 Copy Calling

When comparing predicted and observed values, we use two steps in making the final labeling of copy numbers. First, we identify observed values that are significantly different from predicted values, then we compute the most likely copy state that would generate the significant observed values.

When identifying significant deviations from the predicted values, we consider a sliding window of bins within *k* steps of the query point (we use *k* = 5 in all experiments). For bin location *j*, we consider the set *W* = *{j − k*, …, *j*, …, *j* + *k}*, removing indices that go beyond the bounds of the chromosome. For each location *w* in *W*, we compute the probability of the observed value *o*_*w*_, using a Poisson distribution with the predicted value *p*_*w*_ as a mean.

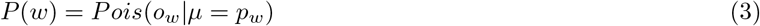

These probabilities are averaged over *W*, and converted into a quantile using an empirical distribution for the mean of random probabilities (known as a Bates distribution[27]). This quantile value gives us a [0, 1] indicator of how extreme the observed values around *j* are compared to the predicted values, with values near 0 or 1 indicating potential CNV loss or gains, respectively. Once a value is computed for all locations across all cells, we use FDR to identify cutoff values for significantly high or low values (*α* = 0.025 for each). These values are deemed as CNV events, and sent to the next step to assign a copy number.

When assigning copy numbers to significant values, we consider the multipliers *{*0.1, 0.5, 1, 1.5, 2, 3*}*. For each multiplier *m*, we look at the likelihood *Pois*(*o*_*w*_|*μ* = *mp*_*w*_). Whichever multiplier gives the highest likelihood is assigned as the copy number, with the special case that the multiplier of 0.1 is assigned the copy number of 0. All non-significant windows are assigned a copy number of 1 (normal). We treat copy numbers less than 1 as losses, and copy numbers greater than 1 as gains. While we stop at copy number 3, higher values could be included for increased resolution of high ploidy cells.

## Acknowledgements

We would like to thank Dr. Andrew Adey and Dr. Hisham Mohammed for their help and feedback in manuscript preparation. We would also like to thank Dr. Ryan Mulqueen for his guidance with other copy caller algorithms.

