## Supplemental Material for "Robust CNV detection using single-cell ATAC-seq"

### Supplemental Materials

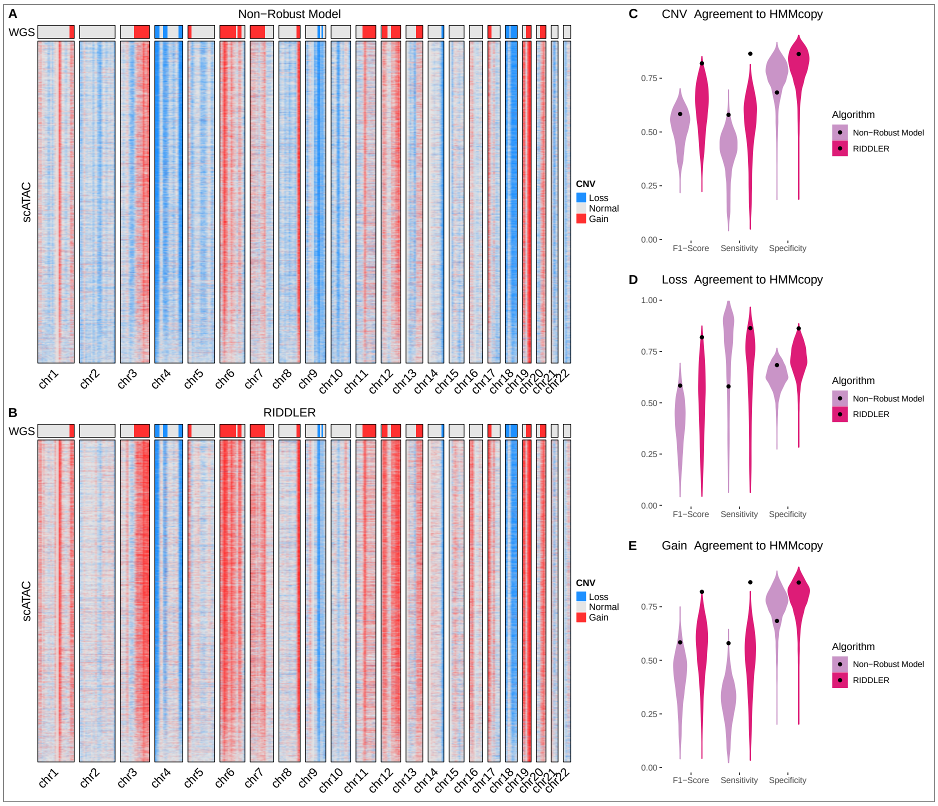


Figure 1: A-B) Heatmaps of CNV calls for each algorithm on SNU601 cell line, with WGS CNVs as reference. C-E) Metrics comparing predicted CNVs to WGS calls. The non-robust model performs significantly worse, classifying less significant gains as normal and large portions of normal genome regions as losses.

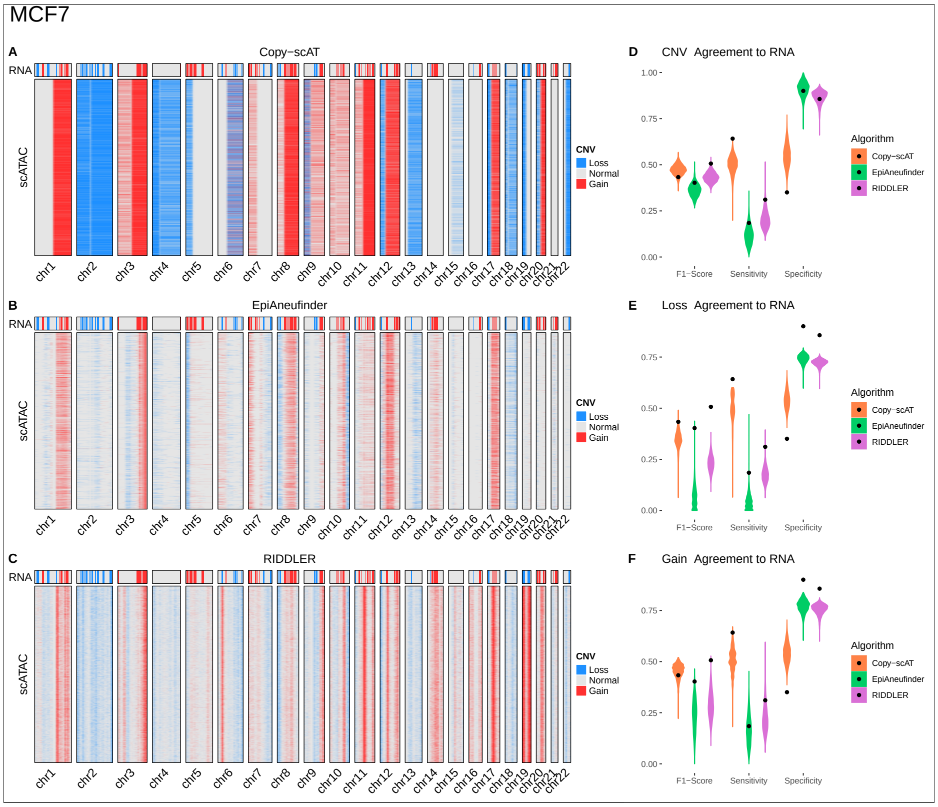


Figure 2: A-C) Heatmaps of CNV calls for each algorithm on MCF7 cell line, with RNA CNVs as reference. D-F) Metrics comparing predicted CNVs to RNA calls. Omitted from main figures for space consideration.

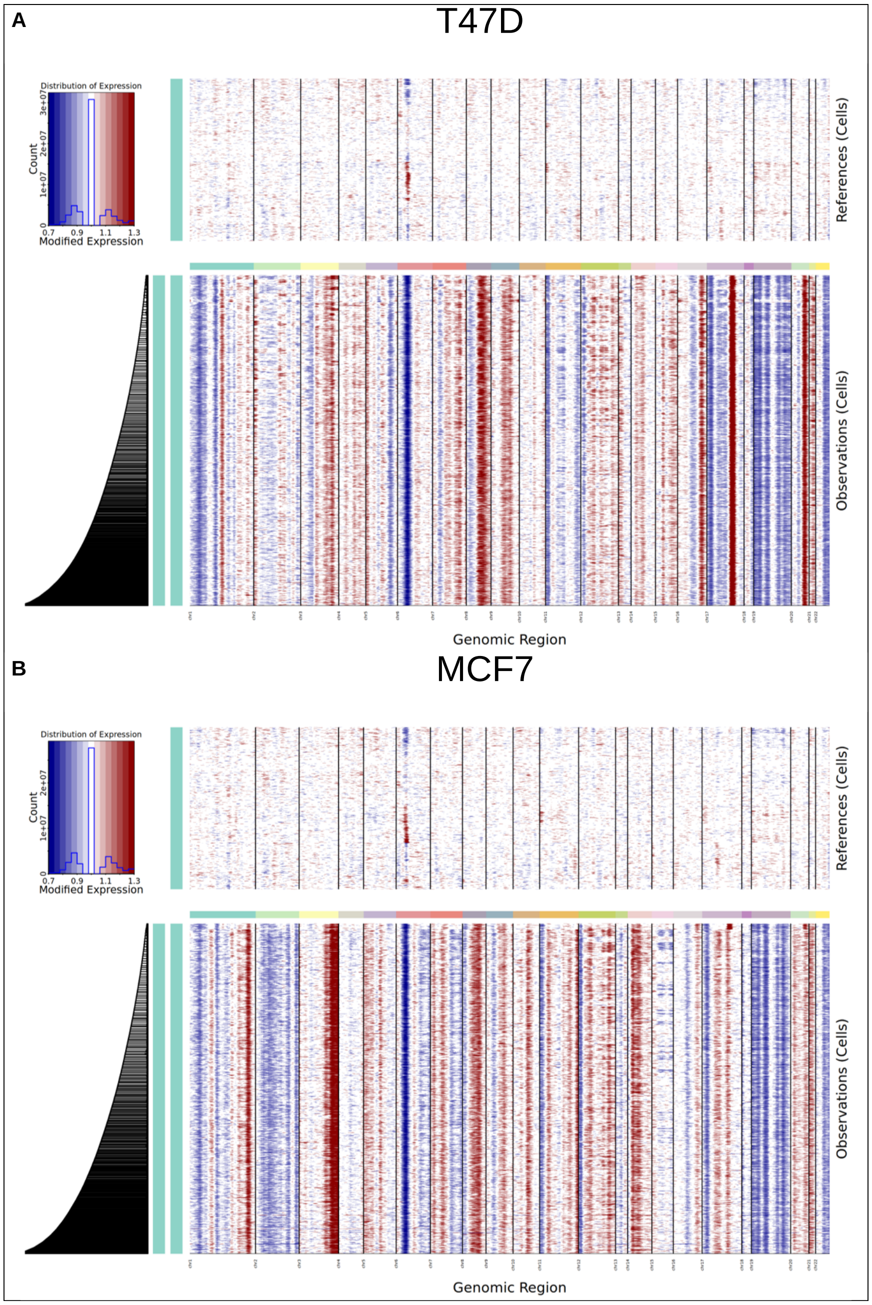


Figure 3: InferCNV output heatmaps for A) T47D and B) MCF7, with PBMS data used as reference for both. Neither cell line showed any indication of CNV substructure.
